# Impact of crowding on the diversity of expanding populations

**DOI:** 10.1101/743534

**Authors:** Carl F. Schreck, Diana Fusco, Yuya Karita, Stephen Martis, Jona Kayser, Marie-Cécilia Duvernoy, Oskar Hallatschek

## Abstract

Crowding effects are key to the self-organization of densely packed cellular assemblies, such as biofilms, solid tumors, and developing tissues. When cells grow and divide they push each other apart, remodeling the structure and extent of the population’s range. It has recently been shown that crowding has a strong impact on the strength of natural selection. However, the impact of crowding on neutral processes remains unclear, which controls the fate of new variants as long as they are rare. Here, we quantify the genetic diversity of expanding microbial colonies and uncover signatures of crowding in the site frequency spectrum. By combining Luria-Delbrück fluctuation tests, lineage tracing in a novel microfluidic incubator, cell-based simulations, and theoretical modeling, we find that the majority of mutations arise behind the expanding frontier, giving rise to clones that are mechanically “pushed out” of the growing region by the proliferating cells in front. These excluded-volume interactions result in a clone size distribution that solely depends on *where* the mutation first arose relative to the front and is characterized by a simple power-law for low-frequency clones. Our model predicts that the distribution only depends on a single parameter, the characteristic growth layer thickness, and hence allows estimation of the mutation rate in a variety of crowded cellular populations. Combined with previous studies on high-frequency mutations, our finding provides a unified picture of the genetic diversity in expanding populations over the whole frequency range and suggests a practical method to assess growth dynamics by sequencing populations across spatial scales.

**Significance Statement:** Growing cell populations become densely packed as cells proliferate and fill space. Crowding prevents spatial mixing of individuals, significantly altering the evolutionary outcome from established results for well-mixed populations. Despite the fundamental differences between spatial and well-mixed populations, little is known about the impact of crowding on genetic diversity. Looking at microbial colonies growing on plates, we show that the allele frequency spectrum is characterized by a simple power law for low frequencies. Using cell-based simulations and microfluidic experiments, we identify the origin of this distribution in the volume-exclusion interactions within the crowded cellular environment, enabling us to extend this findings to a broad range of densely packed populations. This study highlights the importance of cellular crowding for the emergence of rare genetic variants.

## Introduction

Environmental factors often structure the spatial organization of growing cellular populations, such as microbial biofilms^1^, developing embryos and differentiating tissues^2^, as well as solid tumors^3–6^. Advances in lineage tracing techniques are progressively revealing that in many of these cases growth is non-uniform across the population, as it strongly depends on the mechanical and biochemical cues experienced by each cell^4,5,7–14^. Non-uniform growth can favor individuals based on their spatial locations rather than their fitness^6,15–18^ and as such can dramatically impact the evolutionary fate of the population.

The interplay between evolution and growth has been extensively investigated in the context of range expansions, in which populations grow by invading surrounding virgin territory^19–25^. In cellular range expansions, growth is often limited to a thin layer of cells at the expanding front of the population (the *growth layer*) due to processes like nutrient depletion, waste accumulation, mechanical pressure, or quorum sensing in the bulk^26–32^. Recent studies have revealed that this growth constraint generates an excess of high-frequency mutations in microbial colonies^33^ and colorectal cancer xenografts^6^. Remarkably, the size distribution of these large clones is exclusively determined by the surface growth properties of the population through a phenomenon called allele surfing^19,34^.

The distribution of low-frequency mutations, however, remains an open question. Assuming a mutation rate of 10^3^ mutation/genome/generation (typical of microbes) and a population size of 10^8^-10^9^ cells, a total of 10^5^-10^6^ mutations are generated during population growth. Yet, experimentally only approximately 0.001 % of these mutations have been captured by population sequencing in the case of bacterial colonies and tumors^33,35–37^. This suggests that low-frequency mutations constitute the majority of genetic diversity in the population, but since their frequency is often below the detection limit of population sequencing, they go unaccounted for. As a single mutant can be sufficient to drive drug resistance^15^, its quantification is imperative to better understand the emergence of resistant cells after drug treatment. While several groups have recently revealed the dynamics of small clones by multicolor lineage tracing in solid tumors^5,6,12^, a quantitative understanding of the dynamics of low-frequency mutations is still lacking. Here, we address this gap by investigating the dynamics of low-frequency mutations utilizing an expanding microbial colony as a model system.

To probe the low-frequency end of the mutational spectrum, we adapt the classic Luria-Delbrück fluctuation test, normally used to infer mutation rates in well-mixed populations^38^, to microbial colonies. We find that the vast majority of mutations occurring during growth are present at very low frequencies and characterized by a clone size distribution that decays faster than that observed at high frequency^33^. To investigate the origin and statistics of low frequency clones at single-cell resolution in a well-controlled environment, we designed a microfluidic chemostat (the “population machine”) that mimics the growth at the expanding front of a colony. In combination with a newly engineered color-switching *S. cerevisiae* strain, we track clonal lineages for ten generations. Visualization of the clones shows that small clones stem from mutations that occur behind the population’s front. The mutant cells are then pushed towards the bulk of the population by the proliferating cells in front and eventually fall out of the growth layer and stop dividing, limiting the maximum size a clone can reach.

Cell-based simulations show that mechanical cell-cell forces are sufficient to explain the observed low frequency spectrum, and that the spectrum’s behavior is robust to cell-level details such as cell shape and mode of division.

We further develop a theoretical model that captures the essential population genetic process that shapes the low frequency spectrum, extends our results to a broad range of cellular populations,and provides predictions beyond evolutionary neutral populations.

Finally, we discuss a useful sampling strategy to sequence spatially structured populations such as tumors. We show that the spatial position where one takes samples defines which regime of the site frequency spectrum one can capture. Our results suggest that the whole site frequency spectrum can be reconstructed by combining various sampling methods and rescaling.

## Results

### Fluctuation test in bacterial colonies

To assess the clone size distribution of small clones (*<* 10^4^ cells) in *E. coli* colonies grown from single cells to ≈ 10^9^ cells, we adapted the Luria-Delbrück fluctuation test^38^, routinely used to determine spontaneous rates of resistant mutations in well-mixed populations^39–43^, to structured populations like colonies (Fig. 1). Colonies were grown on rich non-selective media, scooped up completely after two days of growth, resuspended, and then plated on selective plates containing nalidixic acid (see Methods). After overnight growth, the selective plates were imaged and the number of resistant colony forming units (CFUs) were counted (Methods).

**Figure 1.**
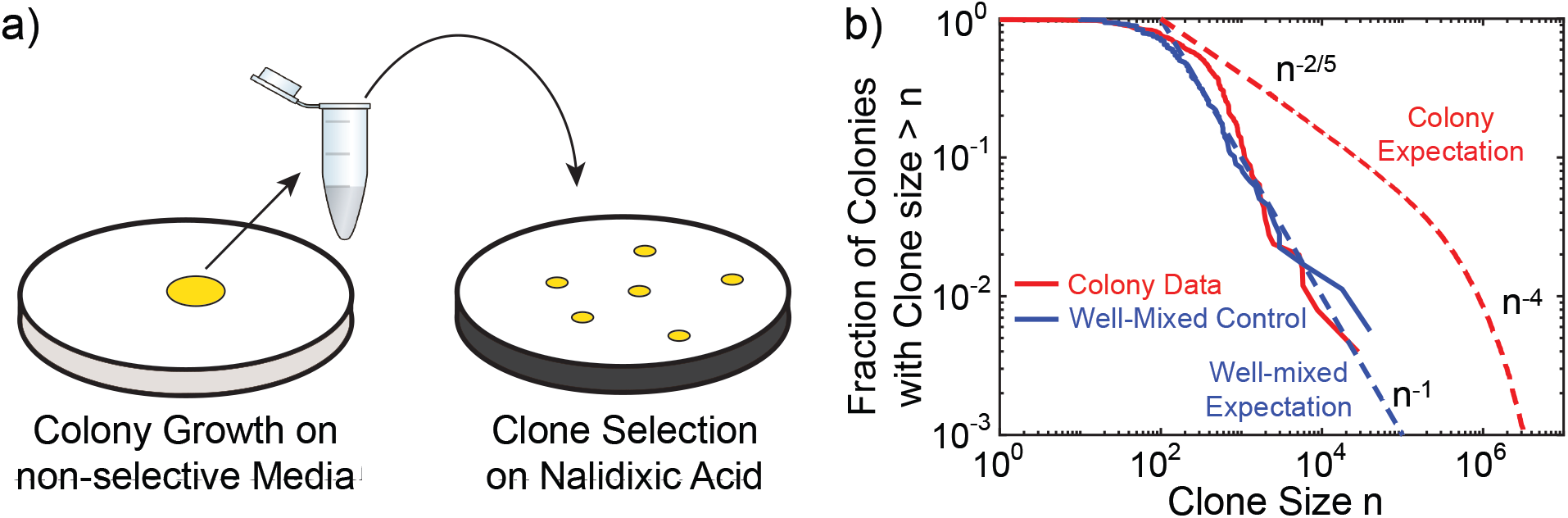
Fluctuation test in bacterial colonies reveals a distinct clone size distribution at low frequencies. (a) Fluctuation test on 234 *E. coli* colonies that were grown for two days, completely harvested and then plated on nalidixic acid. The size of clones corresponding to resistant mutations was determined by counting the number of CFUs on selective plates. (b) Fraction of the sampled colonies carrying at least *n* resistant mutants (red solid line) in comparison with the well-mixed control (blue solid lines). The blue dashed line corresponds to the classic Luria-Delbrück distribution for well-mixed populations (*n*^−1^)^44^, while the red dashed line corresponds to large clones found in colonies (*n*^−2*/*5^ and *n*^−4^ regimes, corresponding to so-called “bubble” and “sector” patterns that were previously characterized^33^.)

The resulting distribution exhibits a decay that resembles the classic Luria-Delbrück distribution typical of well-mixed populations (dashed blue line in Fig. 1), in contrast to the distribution of large mutant clones (*>* 10^5^ cells) previously observed in similar colonies of the same strain via population sequencing (dashed red line)^33^. Indeed, comparison of the clone size distribution pre-factors between colonies and well-mixed populations from sequencing data had previously hinted to the necessary presence of a different distribution regime at very low frequencies^33^. In the following, we investigate the physical origin of these low-frequency clones and characterize their statistics.

### Clone tracking experiments on microfluidics

Because in colonies cell replication is primarily limited to the region near the expanding front, called the “growth layer”^19,45^, most genetic mutations likely occur in this region. In order to visualize the emergence and dynamics of clones over several generations in a well-controlled environment, we designed an *in vitro* growth layer using a microfluidic chamber inoculated with a newly engineered color-switching budding yeast strain (Fig. 2a, b and Methods). In the chamber, whose design is inspired by previous studies^46–50^, all cells grow at the same rate (Fig. S4) and are continuously pushed out as the cells in front proliferate, mimicking the mechanical interactions between cells at the growing edge of a colony in its co-moving frame. By pinning the position of the population front, the device enables tracking the growth layer at single-cell resolution for up to 4 days (Fig. 2a, b and Methods).

**Figure 2.**
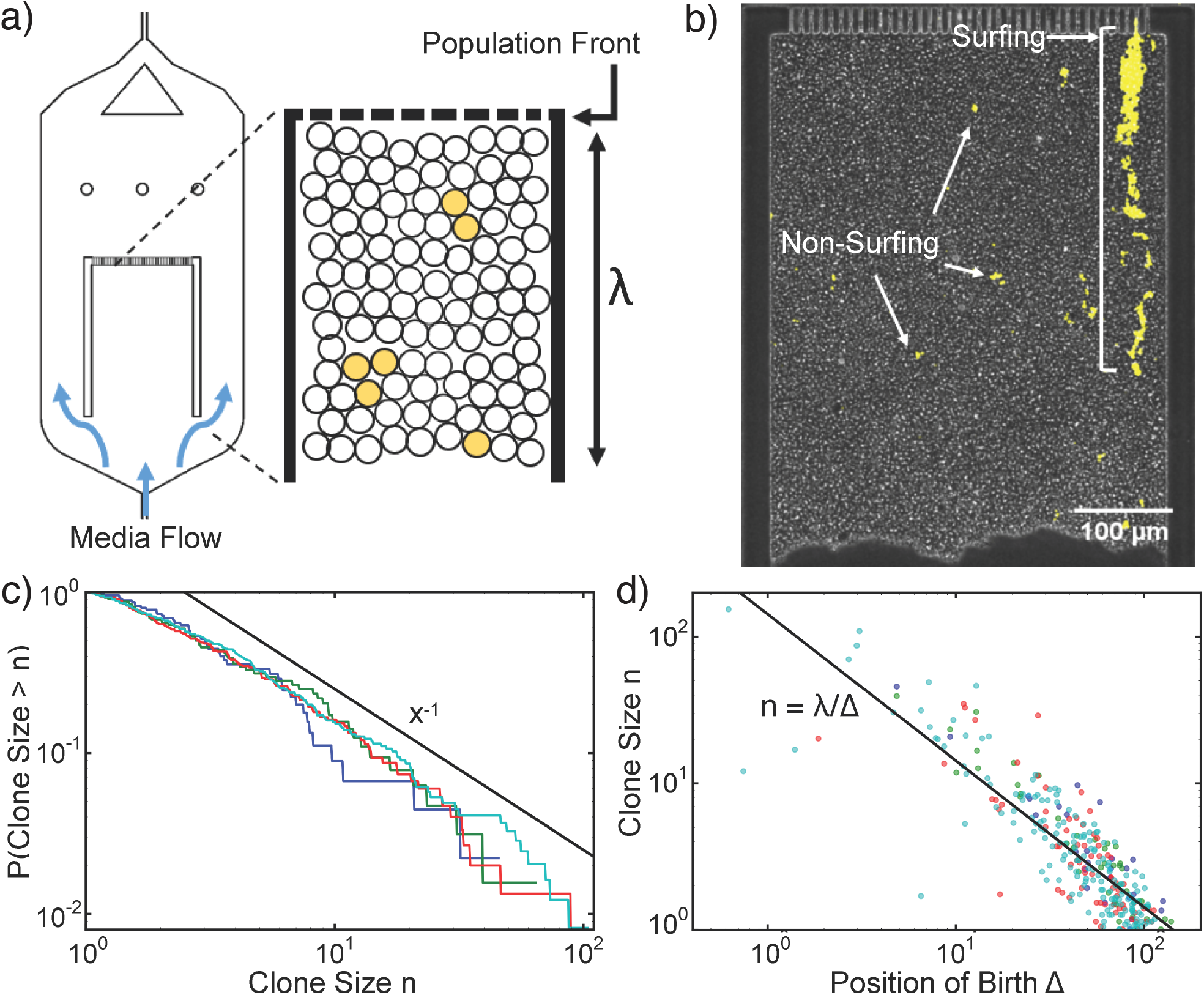
(a-d) Our microfluidic incubator enables the tracking of front dynamics over several generations. (a,b) Schematic and snapshot of microfluidic experiments. Cellular growth within the chamber models the co-moving frame of the growth layer in an expanding colony. Nutrients are supplied from both the top and bottom of the chamber by diffusion so that all cells grow at the uniform rate (Fig. S4). Cells out of the growth layer are flushed away by continuous media flow. (c) Proportion of color-switched cells whose final clone size is greater than *n*, where area is used as a proxy for clone size. The different lines indicate experimental replicas with respectively 45 (blue), 64 (green), 150 (red), 245 (cyan) mutant clones. (d) Relationship between final clone size and distance from the front at which such clone arose. Colors are as in panel (c). The black line corresponds to *λ /* Δ, where *λ* is the size of the chamber and D is the distance from the front.

To quantify the dynamics of clones stemming from a single mutational event, we conducted lineage tracking experiments (Methods and Fig. S3). Since the switch can occur only at cell division, is inheritable and does not measurably change the growth rate (Fig. S5), it effectively behaves like a neutral mutation, whose position and growth can be visually tracked with fluorescent microscopy.

During the course of the experiment, we observed both surfing clones, which are born at the very front, as well as non-surfing clones, which are born behind the front (Fig. 2b). Surfing events, which have been previously investigated^33^, occur rarely and generate very large clones (10^2^-10^3^ cells each) by letting clones stay at the front for some time. By contrast, non-surfing clones cannot reach sizes larger than 100 cells and exhibit completely distinct dynamics. Using clone area as a proxy for size, we obtained the clone size distribution by tracking non-surfing clones for 19-50 hours. The resulting distribution (Fig. 2d) exhibits a power-law decay in agreement with the fluctuation test experiments (Fig. 1b).

The time resolution of this experiment enables us to go beyond the clones’ ensemble behavior, and to track the dynamics the individual clones. Remarkably, we find that clone size is anti-proportional to the birth position of the first mutant (Fig. 2c). This straightforward relationship, despite the complexities of real cellular populations such as cell death, aging of mothers, and feedback of mechanical pressures on growth rate, suggests that a simple physical process may underlie low-frequency clones.

### Mechanical simulations

To gain an intuition into whether the physical growth process alone is sufficient to generate the clone size behavior observed in Fig. 2, we employed 2D mechanical simulations where individually-modeled cells proliferate and repel each other upon contact (see Methods)^31,51^. We introduced an explicit growth layer of finite depth *l* within which cells of width *s* grow exponentially at a uniform rate (Fig. 3a). Beyond the growth layer, cells are considered to be in the *bulk* and stop growing. We represented proliferation via budding to mimic our microfluidic budding yeast experiments (Fig. 2).

**Figure 3.**
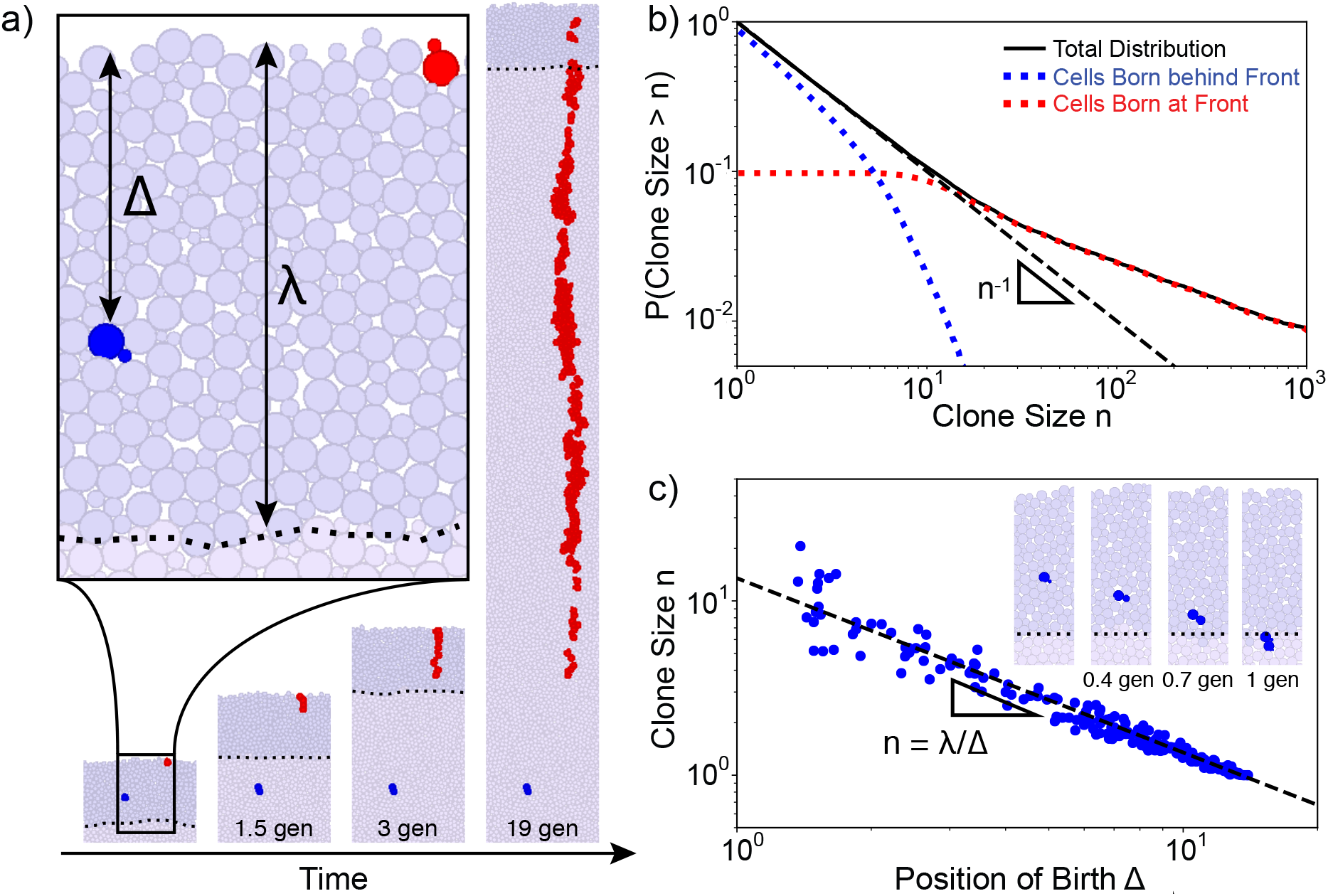
Cell-based simulations show different behaviors between surfing and non-surfing clones. (a) Illustration of the mechanical simulations. Cells lying in the growth layer, defined as the region within a distance Δ *< λ* from the front (dark purple region with dashed line showing back of growth layer), replicate exponentially. In this image, *λ* = 14 cell widths (about 50 μm). As growth proceeds, the front moves at a constant speed and cells behind the front are continuously pushed out of the growth layer by replicating cells in front due to excluded-volume interactions. Mutations can either occur at the very front (red cells) generating a surfing clone, or behind the front (blue cells) generating non-surfing clones that are quickly washed out of the growth layer. Clonal dynamics are shown for the first 20 generations of cellular growth. (b) The full clone distribution (solid black line) can be subdivided in the size distribution of surfing clones (red dotted line), which dominate the high-frequency tail of the distribution, and non-surfing clones (blue dotted line), that dominates the low-frequency behavior. The dashed black line shows the *n*^−1^ prediction. (c) Scatter-plot identifying for each clone (blue dot) the distance from the front at which the mutation first arose and the final clone size upon exiting the growth layer. Surfing clones are by definition clones that arose within 1 cell distance from the front. Non-surfing clones are found to satisfy the relationship *n* = *λ/* Δ, rationalized in Eq. 1 (dashed black line). The inset shows the dynamics of the blue clone a short time (*<* 1 generation) after birth in the reference frame of the front. This clone is born at distance Δ = 7 cells from the front and grows to a size of *n* = 2.

The clone size distribution obtained from simulations exhibits two regimes (Fig. 3b): very small clones (*n* ≲ *λ / σ*, Fig. S1) follow *n*^−1^ while larger clones follow a shallower power-law in quantitative agreement with the allele surfing prediction^33^. Small clones correspond to mutations originating behind the front whereas large clones correspond to mutations originating at the front. When looking at clones arising behind the front, we find that clone size decreases monotonically with the birth position of the first mutant (Fig. 3c).

These results (Fig. 2b,c) agree quantitatively with microfluidic experiments (Fig. 2c,d), showing that the physical process of population expansion is indeed sufficient to generate the *n*^−1^ low-frequency clone distribution. To further investigate whether clone sizes are dependant on cell-level details, we altered the rules of bud site selection in budding cells and also performed simulations of elongated cells (Fig. S7). In both cases, low frequency clones decay as *n*^−1^, suggesting that this underlying phenomena may be described by a simple continuum mathematical model.

### Crowding model of non-surfing clones

To uncover the physical mechanisms underlying non-surfing clones, we developed a mathematical model that describes what we observe in the microfluidic experiments and simulations. As in the simulations, we assumed that the growth rate is uniform within a distance *λ* of the expanding front and zero otherwise. We describe clones in a reference frame that is co-moving with the expanding front, so that rather than accumulating at the edge of the colony, cells are washed out towards the colony bulk (Fig. 3c inset). A mutant of infinitesimal size *δn*_0_ born at a distance Δ from the front will grow until it is pushed out of the growth layer by excluded volume effects from the cells proliferating in front. This happens when the cells in front of the clone have grown to size *λ* to fill the growth layer. Because growth is constant within the growth layer, the mutant will grow by the same relative amount as the layers of cells in front, reaching a final size 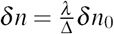. By extending this infinitesimal relation to mutant clones with *n*_0_ = 1 cell at the onset of mutation, we have the prediction (see SI section 1 for finite-size analysis)

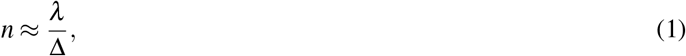

in agreement with cell-based mechanical simulations (Fig. 3b).

Equation 1 translates into a prediction for the clone size distribution when combined with the probability of observing a mutation at distance D. If we assume that the mutation rate is proportional to the growth rate, the probability that a mutation will occur at Δ *< λ* is *P*(Δ)= *λ* ^−1^. Then, the probability of observing a clone of size *n* is

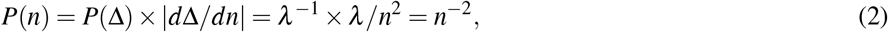

corresponding to a cumulative clone size distribution of *P*(Clone size*> n*)= 1*/n*.

This prediction rests on the assumptions that clone size (*n*) is infinitesimal compared to the growth layer depth (*λ/ σ*) and that cellular growth rate is uniform within the growth layer. We show in SI section 1 that Eq. 1 is robust for finite clones up to *n* = *λ/ σ*, corresponding to mutants born one cell behind the front, which is verified by both microfluidic experiments (Fig. S6) and cell-based simulations (Fig. S1). Additionally, in SI section 2 we show that our prediction also holds in the case of non-uniform growth inside the growth layer, which we verify via simulations in Fig. S9.

### Reconstruction of clone size distribution from subsamples

By characterizing the behavior of low-frequency mutations, a complete picture of the clone size distribution in crowded expanding populations can now be assessed over the entire frequency range. The full distribution (black line in Fig. 4) exhibits three distinct regimes (grey shades in Fig. 4): two regimes for surfing clones that were previously characterized^33^ and one regime for non-surfing clones at low frequencies characterized in this paper. Using random population sequencing, one can capture the complete distribution only by sequencing unrealistically deeply (over 10^5^X coverage). With a typical coverage (10-100X), population sequencing is likely able to assess only the high frequency regimes^33^ (red line in Fig. 4a), and miss the non-surfing bubble behavior that accounts for most of the genetic diversity. However, other sampling strategies can be chosen to take advantage of the spatial proximity of cells that are closely related, a practice that is becoming increasingly frequent in cancer research ^36,37,52–56^. We find that sampling all cells in a small contiguous region of the colony is capable of detecting non-surfing clones (magenta line in Fig. 4a) or the transition between non-surfing and surfing clones (cyan line in Fig. 4a). The data from these contiguous regions can be appropriately rescaled (Fig. 4b, see Methods for rescaling details) in order to recover the complete behavior of the clone size distribution. The local spatial distribution of mutations can therefore be used to identify non-homogeneous growth in the population by sequencing well-chosen sub-samples.

**Figure 4.**
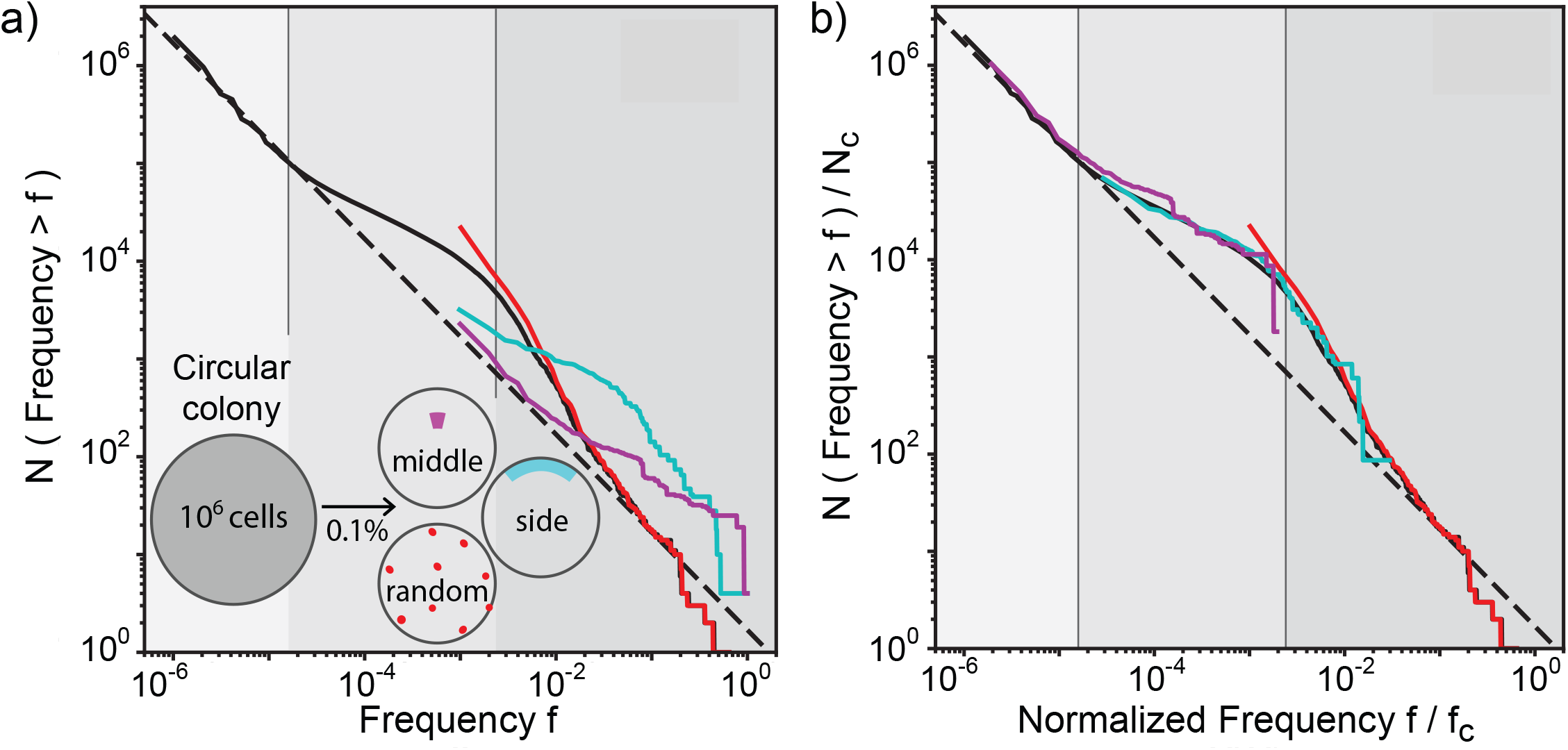
Results from multiple sampling strategies can be combined to infer mutation rate and growth dynamic of the population. (a) Different sampling methods generate distinct clone frequency distributions that highlight distinct properties of the growth dynamics. This in stark contrast with well-mixed populations where the sampling scheme merely affects how well the clone frequency distribution can be resolved. The solid black line shows the clone frequency distribution (clone size divided by population size) of the whole simulated colony (growth layer *λ/ σ* = 14 cells) grown up to 10^6^ cells. We identify three frequency *f* ranges in the site frequency spectrum: (i) for *f <* (*λ/ σ*)*/N*, the distribution is dominated by non-surfing clones; (ii) for (*λ/ σ*)*/N < f <* 0.003, allele surfing dominates generating bubbles and sectors as previously described; (iii) for *f >* 0.003, we see a third behavior, generated by mutations that arise in the first few generations, when the whole microcolony is growing exponentially (*N < π* (*λ σ*)^2^). The grayscale regions correspond to non-surfing bubbles (light gray), surfing bubbles (intermediate gray), and sectors (darkest gray). Sampling 0.1 % of the population (equivalent to a 1000X coverage in sequencing) can target non-surfing small clones and generate their corresponding distribution (middle, magenta), or high-frequency surfing clones (random, red). Sampling an outer segment generates a shifted distribution where distinct trends can be observed. (b) These sampling techniques can be combined to reproduce the entire clone size distribution. The rescaling used here requires only knowledge of the total number of cells in the colony and the size/shape of the sampled region, as are described in SI section 8.

## Discussion

A single resistant cell can seed an entirely new resistant population following an antimicrobial attack. To predict the chances of success of a drug therapy, it is therefore crucial to assess not just the high frequency mutations, but also the rare ones present in small clones after the incubation period. In well-mixed populations, the probability that a mutation carried by at least a frequency *n* is 1*/n* across the entire frequency range. Allele surfing, a hallmark of spatial growth, has been shown to give rise to a different probability distribution characterized by an excess of mutational jackpot events^33^. Here, we have shown that, while allele surfing can explain the behavior of large clones, it fails to describe the majority of mutations which reach much lower frequencies.

Crowded growth in dense populations leads to clones whose final size is determined not by *when*, but *where* a mutation first arose relative to the expanding front. Large surfing clones, which are well described by the surface growth properties of the population, arise at the very front of the expanding edge^45^. However, most mutations occur behind the front, are pushed into the population bulk by proliferating cells near the front, and reach only small final clone sizes. This process leads to a reproducible relationship between final clone size and initial position of the first mutant cell, generating a clone size distribution different from that of surfing clones.

Because clone size is only determined by the relative position to the front, our argument to derive the full distribution is not limited to two-dimensional colonies expanding at the outer edge, but can be applied to a wider class of populations. Theoretical analysis predicts that these results hold in any system where (i) growth rate varies only along the direction of expansion, (ii) a reference frame exists where the growth profile is constant over time, and (iii) the mutation rate per generation is proportional to the growth rate (see SI). Under these conditions, the clone size distribution describing small clones decays like *n*^1^ up to a critical size that depends only on the growth layer depth but is independent of the number of dimensions (circular colonies vs. solid tumors, see SI), origin of growth (outer vs. inner growth, see SI) or mode of proliferation (budding vs. symmetric division, see SI), demonstrating the robustness of the distribution.

Our model allows to extend the theoretical predictions to more complex evolutionary scenarios. For instance, we predict deviations from the *n*^−1^ power law in the cases where mutations confer selective effects (see SI section 4, simulation confirmation in Fig. S8) and where mutation rate and net growth rate are not proportional as may be the case during necrosis (see SI section 4).

As the population expands, the majority of mutations are left behind in the bulk, forming a reservoir of genetic diversity in the population. In a typical microbial colony with a growth layer of approximately 100 cells, these mutations would account for more than 99% of the genetic diversity. Analogously, in a solid tumor, they would be responsible for the vast majority of the intra-tumor heterogeneity, while being largely undetectable by population sequencing. As this class of mutations is the most abundant, it is likely to harbor those rare mutations that can confer resistance. Upon environmental changes that kill the surrounding wild-type, these mutants can be spatially released and thus rescue the population from extinction^33,57^. It is therefore crucial to develop methodologies that enable their detection.

In well-mixed populations, the detection power is limited by the sequencing coverage one can afford. Still, because the clone size distribution is characterized by a single process across the full range of frequencies, it is possible to estimate mutation rates and selection effects using a reasonable depth of sequencing. Here we have shown that this procedure cannot be applied to crowded populations growing in space, since the shape of the clone size distribution is controlled by very different processes at low and high frequencies. A way around this problem consists in exploiting the spatial arrangement of the population. Neighboring cells are likely to be more closely related than cells farther apart, therefore concentrating sampling power to one or few locations in the population would allow to reach deeper into the low-frequency regime and measure important population genetic parameters like the mutation rate.

In the context of cancer, where there are active debates on how to distinguish selection from neutral evolution^58,59^, our findings highlight the additional challenge of distinguishing selection effects from non-uniform growth that is exclusively driven by spatial constraints. Recent work has recognized similar effects in experiments and simulations, proposing phenomenological models of the tumorogenic evolutionary process^5,12,37,60^. Here we offer a microscopic, physical model of evolutionary dynamics that is consistent with the patterns of genetic diversity in solid tumors (*n*^1^ distribution in^5,37^) and which is flexible enough to provide insight into the effects that different evolutionary and demographic processes have on the statistics of rare mutants. By taking advantage of the spatial proximity of closely-related cells, this model offers rational sampling strategies for probing clone size distributions that can be useful for characterizing intra-tumor heterogeneity in cancer research^52–56^. These results can better characterize the growth dynamics of the tumor, which can be used to more precisely identify signatures of selection.

## Method

### Fluctuation test in *E. coli*

The mutator strain *mutT* of the bacterium *E. coli* was used for the fluctuation test experiment on nalidixic acid. The spontaneous mutation rate in this strain was estimated to be approximately 2 10^7^ per generation from the fluctuation test in the well-mixed control, which is consistent with previously reported values^61^. Colonies starting from single cells were grown on plates with LB and 2 % agar at 37°C for 30 hours up to a population size between 10^8^ and 5 10^8^ cells. Each of the 234 colonies was completely scooped from the plate with a pipette tip and resuspended in PBS. A 100X dilution of the resuspension was stored in the fridge for further analysis, while the rest was plated on selective plates containing LB, 2 % agar and 30 μg/mL of nalidixic acid for CFU count. The selective plates were incubated overnight at 37°C and imaged the following day. CFU count was determined semi-manually with a built-in ImageJ function (see below). If selective plates exhibited more than 400 CFUs, the set-aside 100X dilution was itself plated on nalidixic acid, incubated overnight, and imaged the following day to better estimate the size of large mutations. In the control experiment under well-mixed condition, populations were started from about 50 cells in 200 μL of LB and incubated on a table-top shaker overnight up to saturation. The final population size was estimated to be between 10^8^ and 10^9^. Each of the 178 well-mixed populations was treated similarly as described above.

### Colony counting on plates

Images of colonies on plates were thresholded and binarized using ImageJ. Thresholding was done manually for each image to minimize the effect of noise, such as dust particles, smudges, or glares. Colonies near the rim of the plates were excluded to avoid an edge effect. Colony counting was done automatically with the Analyze particles function of ImageJ. The final clone size of the well-mixed populations control was rescaled by 10 to take into account the different final population size and to better visualize the comparison with the data from colonies.

### Mechanical simulations

Cells are modeled as 2D rigidly-attached disks of width *σ* that proliferate via budding. Upon division, cells divide in polarly, with newly-formed buds retaining the orientation of their mothers. Cells interact with each other upon contact via purely repulsive elastic forces and move via overdamped Stokesian dynamics^31^. To mimic diffusion of nutrients into the population from the exterior, we allow only cells within a distance *λ* from the front to actively grow while the rest of the population remains in stationary phase. In order to simulate a flat geometry, we impose periodic boundary conditions in the horizontal direction so that the populations expands outward only in the vertical direction. To calculate the frequency of neutral mutations, we periodically label 40, 000 newly-born cells and track their descendants.

### Fabrication of microfluidics

The microfluidics was fabricated by soft lithography^62^. The master mold was made by spin-coating (CEE 100 spin coater, Brewer Science) a 10 μm - thick layer of negative photoresist (SU8-2010, MicroChem) on a silicon wafer (WaferNet). The photoresist was patterned by photolithography on a mask aligner (Hybralign 200, OAI) through a chrome photomask (Compugraphics). The thickness of the pattern was measured by a stylus meter (Dektak3030, Bruker). Polydimethylsiloxane (PDMS, Sylgard 184, Dow Corning) was mixed with the crosslinker in 10-to-1 ratio and poured on the mold. After being cured at 60 C overnight, the PDMS was peeled off from the mold and punched holes in for inlets and outlets. The chip was bonded to a glass coverslip after O_2_ plasma treatment by a reactive ion etcher (Plasma Equipment Technical Services). Prior to cell culture, 0.1 % bovine serum albumin (Sigma-Aldrich) was loaded into the device to reduce the interaction between cells and the substrate.

### Yeast Strain

The microfluidics experiments were conducted with the *S. cerevisiae* strain yJK10, derived from strain yDM117 (courtesy of Jasper Rine, University of California, Berkeley). yJK10 employs a Cre-loxP recombination system to switch stochastically from a red (yEmRFP) to a green (yEGFP) fluorescent state, as previously published^33,63^. Using an estradiol-inducible Cre construct allowed us to optimize the average switching rate for our experiments^64^. For all experiments, we used a concentration of 1.6 nM *β* -estradiol corresponding to a switching rate of 7.1 ± 4.8 × 10^−4^ per cell per generation (estimated from the number of observed switching during the microfluidics experiments). In principle, the relative fitness between switched and unswitched cells can be set via the differing cycloheximide susceptibility of both states. However, while we did not perform any variation of relative fitness in this study, we chose to use yJK10 to maximize comparability of our results to ongoing and future investigations involving this strain. Under our experimental condition, the relative fitness between the two states (*s* = 0.022 ± 0.040) is sufficiently small to be neglected (Fig. S5). See the SI section 4 for the effect of non-zero *s* on the power-law exponent of the distribution of clone size.

### Clone tracking in microfluidics

The microfluidic growth chamber was designed as a population version of the mother machine^50^. A suspension of yJK10 cells in an exponential phase was injected into the device with YPD culture medium. After overnight culture, cells grew and filled up the growth chamber. At this point, 1.6 nM *β* -estradiol was added to the culture medium to induce color switching (the switching rate was about 10^−3^ per cell division). Subsequent growth was imaged using time-lapse microscopy on an inverted microscope (IX81, Olympus) with a 10X objective every 10 minutes for 2-4 days. The taken GFP images (color of switched cells) were binarized by Otsu’s method^65^, and the dynamics of the clones were manually tracked on Matlab (Mathworks) and ImageJ (NIH). Throughout the experiment, the temperature was controlled at 30 C by a microscope incubator (H201-T, Okolab), and the flow rate of the medium was regulated by syringe pumps (neMESYS, CETONI) at 15 μL/h. The growth rate of cells was uniform across the chamber under our experimental condition (Fig. S4)^66^.

## Data availability

All data, codes, scripts and CAD files for microfluidics are available upon request.

## Acknowledgement

Research reported in this publication was supported by the National Institute of General Medical Sciences of the National Institutes of Health under award R01GM115851, a National Science Foundation CAREER Award (#1555330), a Simons Investigator award from the Simons Foundation (#327934), and the National Energy Research Scientific Computing Center, a US Department of Energy Office of Science User Facility operated under contract number DE-AC02-05CH11231.

## Data availability

All data generated and analyzed in this study are available upon request.

## Code availability

All codes used for the data analysis and the simulations in this study are available upon request.

