## Supplementary Materials for "Impact of crowding on the diversity of expanding populations"

### Supplementary Information

#### 1 Finite size effects in minimal model of clone size distribution

Here, using a 1D mathematical model of growth layer expansion, we derive a relationship between birth position ($\Delta$) and clone size ($n$) without relying on an assumption that clone sizes are infinitesimal.

We first consider an infinitesimally small mutant of width $d\sigma_0$ born at a distance $\Delta$ from the front. This mutant will grow until it is pushed out of the growth layer by the cells proliferating in front of it, which occurs when the thickness $\Delta$ of cells in front has reached width $\lambda$ filling the growth layer. Because growth is constant within the growth layer, the infinitesimal mutant will grow by the same relative amount as the thickness of cells in front, reaching a final width $d\sigma_f = \frac{\lambda}{\Delta} d\sigma_0$.

Next, we consider a mutant with an initial finite width $\sigma_0$ centered at a distance $\Delta$ from the front by subdividing it into many infinitesimal mutant segments over the range $[\Delta - \sigma_0/2, \Delta + \sigma_0/2]$. Since each infinitesimal segment $d\Delta'$ satisfies the relationship above, the final size of the mutant clone will be

$$\sigma_f = \int_{\Delta-\sigma_0/2}^{\Delta+\sigma_0/2} \frac{\lambda}{\Delta'} d\Delta' = \lambda \log\left(\frac{\Delta+\sigma_0/2}{\Delta-\sigma_0/2}\right). \tag{3}$$

Assuming that the initial width $\sigma_0$ corresponds to one cell width, we refer to $n = \sigma_f/\sigma_0$ as the final clonal size and express $\lambda$

$$n = \lambda \log\left(\frac{\Delta+\sigma_0/2}{\Delta-\sigma_0/2}\right). \tag{4}$$

In the limit where $\Delta \gg 1/2$ (or equivalently $n \ll \lambda$)

$$n \approx \frac{\lambda}{\Delta/\sigma_0}, \tag{5}$$

This relationship between position at birth and final clone size is consistent with what we predict in the main text for infinitesimal clones (Eq. 1) and find in cell-based simulations (Fig. 3b).

This approximation underestimates the final clone size, with the largest errors corresponding to mutations that occur closest to the front ($\Delta \ll \lambda$). However, the approximation works very well even for the largest possibly non-surfing clones that originate at $\Delta = \sigma_0$, corresponding to a relative error of $|(n_{\text{exact}} - n_{\text{approx}})/n_{\text{exact}}| = 0.1$. Clones born closer to the front ($\Delta < \sigma_0$) tend to surf (Fig. 3c), leading to a qualitative change in the clone size distribution that is highly dependent on the granular nature of the cell colony.

The relationship between position at birth and final clone size, which holds for each clone individually, translates into a prediction for the global clone size distribution when combined with the probability of observing a mutation at distance $\Delta$. We assume that the mutation rate is proportional to the growth rate (no death), meaning that mutations occur with a certain probability only when a new cell is born. Since growth is constant within the growth layer, the probability that a mutation will occur at $\Delta$ (for $0 < \Delta < \lambda$) is $P(\Delta) = \lambda^{-1}$. By inverting Eq. 4 in order to calculate $d\Delta/dn$

$$\Delta = \frac{1}{2}\frac{e^{n/\lambda}+1}{e^{n/\lambda}-1}, \tag{6}$$

we can obtain the probability of observing a clone of size $n$

$$P(n) = P(\Delta)\left|\frac{d\Delta}{dn}\right| = \frac{e^{n/\lambda}}{\lambda^2(e^{n/\lambda}-1)^2}, \tag{7}$$

If $n \ll \lambda/\sigma_0$, we find the approximate relation we had before $P(n) \approx n^{-2}$ (and cumulative distribution $P(n > \text{Clone size}) \approx n^{-1}$).

#### 2 Extension to non-uniform growth layer

We built an ODE model able to explain the form of the clone size distribution we observe for an arbitrary one-dimensional growth profile $k(z)$. At time $t = 0$, a mutant cell is born a distance $\Delta$ behind the front. We assume that the growth rate $k(z)$ depends only on the $z$ position of the cell measured as distance from the front in units of cell widths and that $k(z)$ is constant over the length of one cell. The clone size $n$ will evolve in a position

$$\dot{n} = \int_{z(t)}^{z(t)+n} k(z')dz' \approx k(z(t))n$$

where $z(t)$ is mutant clone's position at time $t$. We have taken the limit that the characteristic clone size is smaller than the lengthscale on which $k$ decays. Formally, the final size of the clone will be:

$$n_\infty = n_0 \exp\left[\int_0^\infty k(z(t))dt\right]$$

where $n_0$ is the initial size of the clone (in most cases 1) The element's position, $z$, will move away from the front with velocity:

$$\dot{z} = \int_0^z k(z')dz'$$

if we choose a frame of reference in which the front is 'pinned' at $z = 0$ while the bubble is pushed back along the nutrient profile. A key assumption here is that the growth profile is stable in relation to the front. Now, we can write the asymptotic clone size as:

$$n_\infty = n_0 \exp\left[\int_\Delta^\infty \frac{k(z)}{\dot{z}}dz\right] = n_0 \exp\left[\int_\Delta^\infty \frac{k(z)}{\int_0^z k(z')dz'}dz\right]$$

Note that in order for the asymptotic area to be well-defined, the integral over $z$ must be finite. We make the change of variables:

$$\begin{aligned} \kappa_z &\equiv \int_0^z k(z')dz' \\ d\kappa_z &= k(z)dz \end{aligned}$$

Therefore:

$$n = n_0 \frac{\kappa_\infty}{\kappa_\Delta}$$

where we have dropped the subscript on $n$. We see that $\kappa_\infty$ must be a finite constant to have a finite asymptotic area, so this further constrains our choice of $k(z)$.

The clone size distribution $P(n)$ can be written as

$$P(n) = \left|\frac{da}{d\Delta}\right|^{-1} P(\Delta),$$

and from the relationship above we have that

$$\left|\frac{dn}{d\Delta}\right| = n_0 \frac{\kappa_\infty}{\kappa_\Delta^2} k(\Delta)$$

If we assume that the probability of mutating is proportional to the growth rate, then $P(\Delta) \sim k(\Delta)$. It thus follows that

$$P(n) \sim \frac{\kappa_\Delta^2}{n_0 \kappa_\infty} \sim \frac{1}{n^2}.$$

The result holds for exponential growth profiles, power law profile with small and large $z$ cutoffs, and Monod type profiles of the form $\frac{e^{-z}}{1+e^{-z}}$. We explicitly test this prediction for mechanical cell-based simulations with exponential profiles in Fig. S9.

Another interesting aspect of this analysis is that the final bubble area depends on its position at birth through the term $\kappa_\Delta$. We see that $\kappa_\Delta$ is a measure of the total amount of available biomass between the bubble's birth position and the front. In other words, the bubble size is dictated by global properties of the nutrient profile.

#### 3 Selection: single effect size

If the mutations are not neutral, the mutant population will grow at a different rate compared to the WT. We assume that this difference is given by a multiplicative constant, so that if the WT grows according to $k(z)$, the mutant grows according to $(1+s)k(z)$ where $s > -1$ is the "fitness difference" between the two strains.

Using the same analytical derivation as in the previous section, we find that

$$n = n_0 \left(\frac{\kappa_\infty}{\kappa_\Delta}\right)^{1+s}.$$

Note that for neutral mutations ($s = 0$) we recover the old result. We now get for the probability distribution (conditioned on $s$):

$$P(n|s) \sim n^{-\frac{2+s}{1+s}}$$

corresponding to cumulative distribution $P(\text{Clone size} > n) \sim n^{-1/(1+s)}$. See Fig. S8 for verification of this prediction in mechanical cell-based simulations.

This prediction follows our intuition: if $s < 0$ (deleterious mutations) the bubble distribution falls off more steeply, whereas it becomes more broad as we get to larger positive fitness effects.

#### 4 Selection: distribution of fitness effects

A distribution of fitness effects will also create noticeable distortions in the clone size distribution. For small $s$, we have:

$$P(a|s) \sim n^{-2+s}$$

For a distribution of fitness effects $P(s)$, we have the clone size distribution:

$$P(n) = \int P(n|s)P(s)ds \sim n^{-2}\langle n^s \rangle_s$$

where $\langle n^s \rangle_s$ is related to the generating function of the distribution of fitness effects, $\langle e^{zs} \rangle_s$, evaluated at $z = \log n$.

Let's assume $s \sim \mathcal{N}(0,\sigma)$, so we have:

$$P(n) \sim n^{-2} \int n^s e^{s^2/\sigma^2} ds$$

where we have dropped any constant factors. We can complete the square and perform the integral to get:

$$P(n) \sim n^{-2} \exp\left(-\frac{\sigma^2 \log^2 n}{4}\right) \approx n^{-2}\left(1 - \frac{\sigma^2 \log^2 n}{4}\right)$$

since $\sigma$ is assumed to be small.

#### 5 Clone size in microfluidic lineage tracking

In microfluidic experiments, we measure the size of clones within a culture chamber at each time point and use the maximum value as a proxy of final clone size. We show here that this approximation does not affect the predicted power-law of the site frequency spectrum.

If a cell is born at distance $\Delta$ from the front with initial length $\sigma_0$, then the initial position of the leading edge of the clone is $z = \Delta + \sigma_0/2$. When the leading edge makes contact with the back of the growth layer ($z = \lambda$), the entire clone is stretched to size $\sigma_f = \sigma_0\lambda/(\Delta + \sigma_0/2)$. Inverting this relationship gives $\Delta = \lambda/n - 1/2$, where the length scale is rescaled by $\sigma_0$. The result slightly deviates from Eq. 1, but $P(n) \propto d\Delta/dn \propto n^{-2}$ holds in this case as well.

#### 6 Clone size distribution with non-homogeneous death rate

The clone size distribution for non-surfing clones behaves like $n^{-2}$ when assuming that the probability $P(\Delta)$ for a mutation to appear at position $\Delta$ is proportional to the net growth rate $k(\Delta)$ in such position. However, this assumption might break under certain conditions, for instance if a non-homogeneous death rate is present. In the general case, it still holds that the probability of observing a mutant of size $n$ is

$$P(n) = P(\Delta)\left|\frac{dn}{d\Delta}\right|^{-1}$$

and the relationship between final size $n$ and position at birth $\Delta$ remains

$$n \propto \left[\int_0^{\Delta} k(z)dz\right]^{-1}$$

where $k(z)$ is the net growth rate at position $z$. However, further simplifications cannot be made, leading to the general expression

$$P(n) \propto \frac{P[\Delta(n)]}{n^2 k[\Delta(n)]},$$

where the notation $\Delta(n)$ highlights that $\Delta$ is a function of $n$. The functional form of the clone size distribution $P(n)$ will then in general depend on the specific form of $P(\Delta)$ and $k(\Delta)$.

For illustration, we report here an example in which we define the net growth rate $k(z) = \exp(-z/\lambda) = \alpha(z) - \beta(z)$, where $\alpha(z)$ represents the birth rate and is proportional to the mutation rate $P(z)$, while $\beta(z)$ is the death rate. In this case, $n = [1 - e^{-\Delta/\lambda}]^{-1}$ and

$$P(n) \propto \frac{\alpha(\Delta)}{n^2(1-1/n)} = \frac{\alpha(\Delta)}{n(n-1)}.$$

If $\alpha(z)$ is uniform along $z$ and $\beta(z) = 1 - e^{-z/\lambda}$, then the clone size distribution $P(n) \propto \frac{1}{n(n-1)}$, which tends to $n^{-2}$ for large $n$, but deviates from it at small $n$. This would correspond to the case in which replication rate is not affected by position, but death rate increases as we move deeper inside the colony, for instance because of the accumulation of toxic waste.

#### 7 Inferring mutation rate from non-surfing clones distribution and localized deep sequencing

We have shown that for non-surfing clones, the probability that a mutation is larger than size $n$ is $P(\text{Clone size} > n) = 1/n$. Indeed, by definition, a mutation has to be carried by at least $n = 1$ cells and $P(\text{Clone size} > 1) = 1$ as intuition suggests. A related quantity to $P(\text{Clone size} > n)$ that can be observed experimentally is the number of mutations $M(\text{Clone size} > x)$ that are carried by at least a frequency $x$ of the population (Fig. 4). Because the total number of mutations in a population of final size $N$ is approximately $\mu N$, where $\mu$ is the mutation rate per replication, it follows that $M(\text{Clone size} > x) = \mu N P(\text{Clone size} > n) = \mu N/n = \mu/x$. It is therefore possible to estimate the mutation rate $\mu$ from the prefactor of the non-surfing clone regime of $M(\text{Clone size} > x)$ (low-frequency range of the black line in Fig. 4).

If the population is too large or sequencing coverage is too low to observe the non-surfing clone regime, we find that localized bulk deep sequencing can be used (cyan line in fig. 4). In this case, the observed frequency $\hat{x}$ of a mutation represents the frequency in the sequenced sub-population $\hat{N} < N$. However, also the total number of mutations in the sub-population scales like $\mu\hat{N}$. As a result, the observed number of mutations above an observed frequency, $\hat{M}(\text{Clone size} > \hat{x}) = \frac{\mu\hat{N}}{\hat{x}\hat{N}} = \mu/\hat{x}$. Therefore, the prefactor of the power-law can again be used to estimate the mutation rate of the population, even if only part of the population is sequenced.

#### 8 Rescaling of entire colony frequency spectra

In order to rescale the clone frequency distributions from sub-sampled regions, we calculate a characteristic frequency $f_c$ and corresponding value of the cumulative distribution $N_c$. The frequency $f_c$ can be thought of as the frequency that a mutation carried by a single cell in the sub-sample would have in the entire colony.

For the side sampling technique, $f_c$ is determined the the solid angle $\theta$ =(sampled width)/(colony radius) that is inscribed by the sampled region: $f_c = \theta$. For the middle sampled regions, $n_c$ further takes into account the ratio $r$ =(number of cells in fictitious inner colony with radius equal to outer extent of sampled region)/(number of cells in entire colony): $n_c = r\theta$.

$N_c$ is then determined by aligning the smallest of value of $N$ is the subsampled region with with predicted trend $N/N_c = n_c/f$.

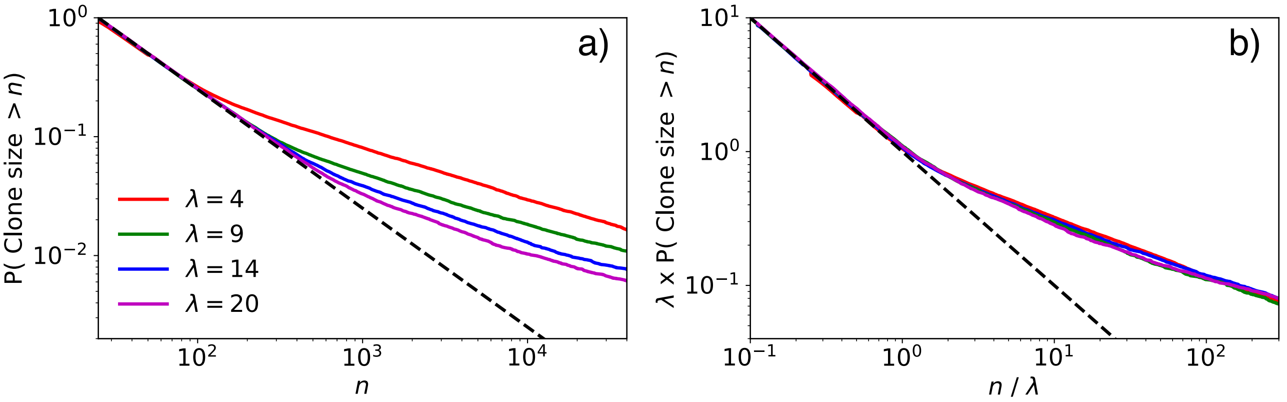


**Figure S1.** (a) Clone size distribution for a range of growth layers depths: $\lambda = 4$, $\lambda = 9$, $\lambda = 14$, and $\lambda = 20$ (units of cell widths). The dashed line shows the $n^{-1}$ prediction. (b) Clone size distribution rescaled by $\lambda$ shows that the $n^{-1}$ regime extends over the range $n = 1$ to $n = \lambda$. For $n > \lambda$, the clone size distribution is dominated by surfing bubbles (Fig. 3a).

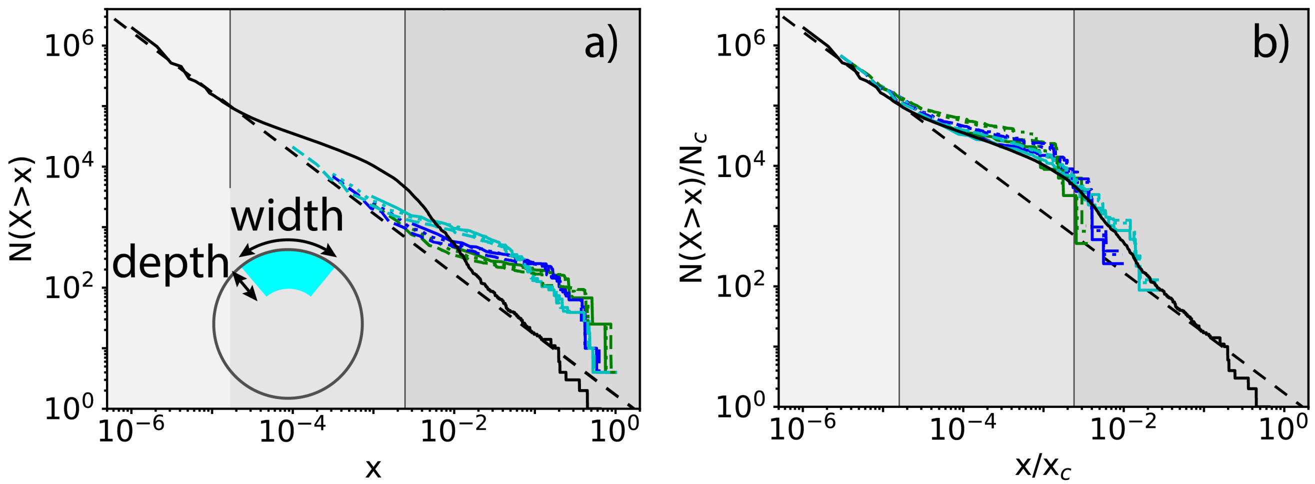


**Figure S2.** (a) Clone size distribution (black) for colony with $\lambda = 14$ cells and radius $R = 602$ cells (total number of cells in colony $= 10^6$). Colored lines show distributions obtained via sub-sampling using side technique with widths 11 cells (green), 36 cells (blue), 112 cells (cyan) and depths of 11 cells (solid line), 36 cells (dashed line), 120 cells (dotted line). Shaded regions correspond to non-surfing bubbles, surfing bubbles, and established sectors. The grayscale regions correspond to non-surfing bubbles (light gray), surfing bubbles (intermediate gray), and sectors (darkest gray). (b) Rescaled distributions, $x_c$ and $N_c$ are described in Section 8.

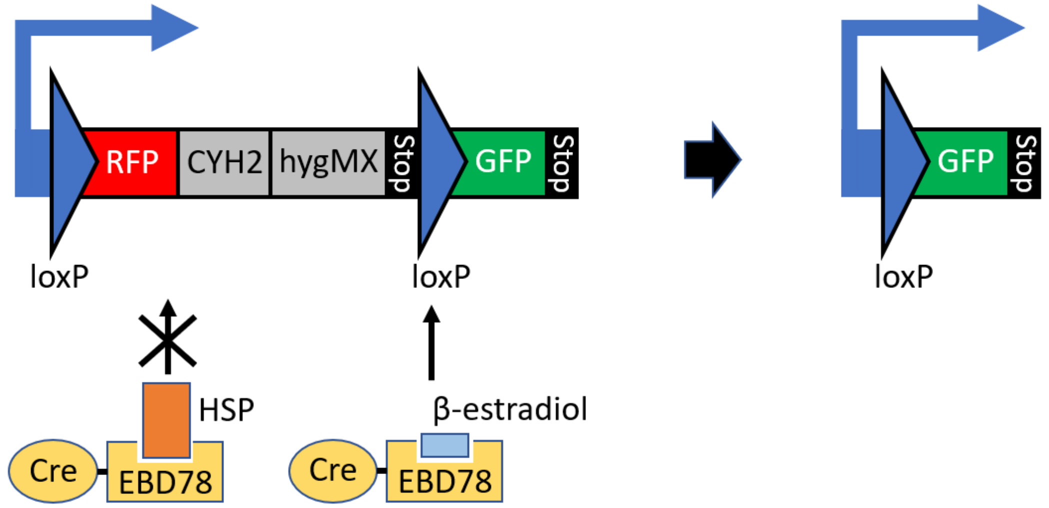


**Figure S3.** A schematic of the engineered *S. cerevisiae* strain yJK10 that stochastically switches the color from RFP (yEmRFP) to GFP (yEGFP). The switching rate is tunable with $\beta$-estradiol, and the fitness advantage/disadvantage of switched cells can be tuned with drugs, hygromycin B and cycloheximide due to an additional cycloheximide resistance allele cyh2Δ::cyh2r logie1995. The genotype of the strain is as follows:
*W303 MATa cyh2Δ::cyh2-Q37E-cs hmlα2Δ::R hoΔ::prSCW11-cre-EBD78-natMX ura3Δ::prGPD-loxP-yEmRFP-tCYC1-CYH2-hygMX-loxP-yEGFP-tADH3*.

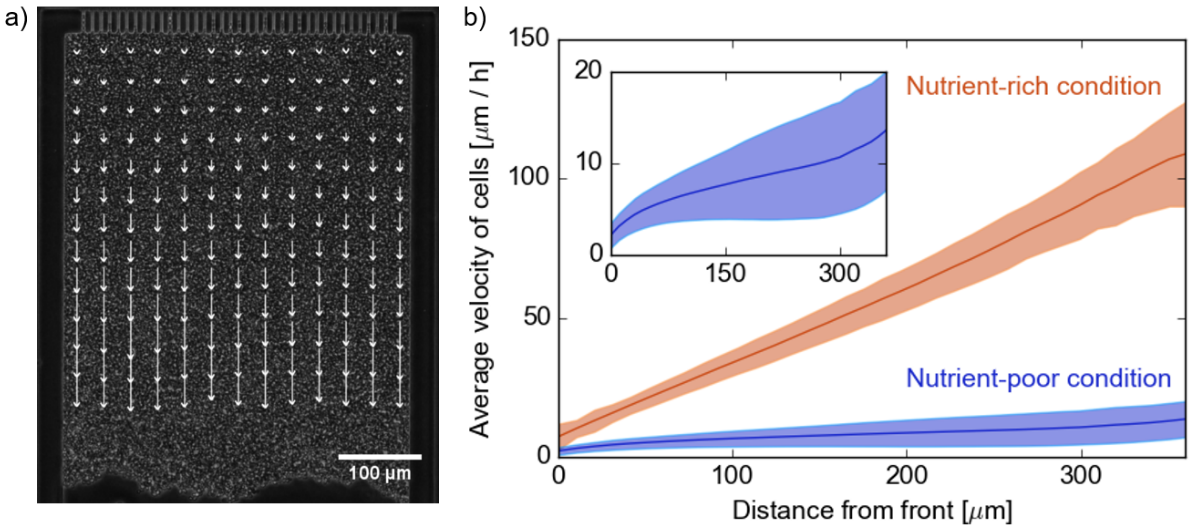


**Figure S4.** (a) A snapshot from the particle image velocimetry analysis Thielicke2014,Thielicke2014-1. Each arrow shows the parallel component of the displacement of a 20x20 μm$^2$ region (32x32-pixel) during one time frame (10 minutes). For the sake of visibility, the length of arrows is rescaled by factor of 2, and the number of arrows is reduced from 37x37 to 13x13. (b) Under our usual experimental conditions (nutrient-rich condition, 2%-glucose YPD media), the velocity field is linear along the growth direction, showing that all cells grow at the same rate. To make sure this method would capture a drop in growth rate, we replicated this experiment under nutrient-poor condition (0.01%-glucose YPD media). As expected, the overall velocity is reduced (slower growth rate) and heterogeneous along the growth direction (see inset), showing a slow down in the middle of the chamber due to nutrient depletion. The error shows the standard deviation of the statistics across horizontal positions and over 100 (nutrient-rich) and 140 (nutrient-poor) time points.

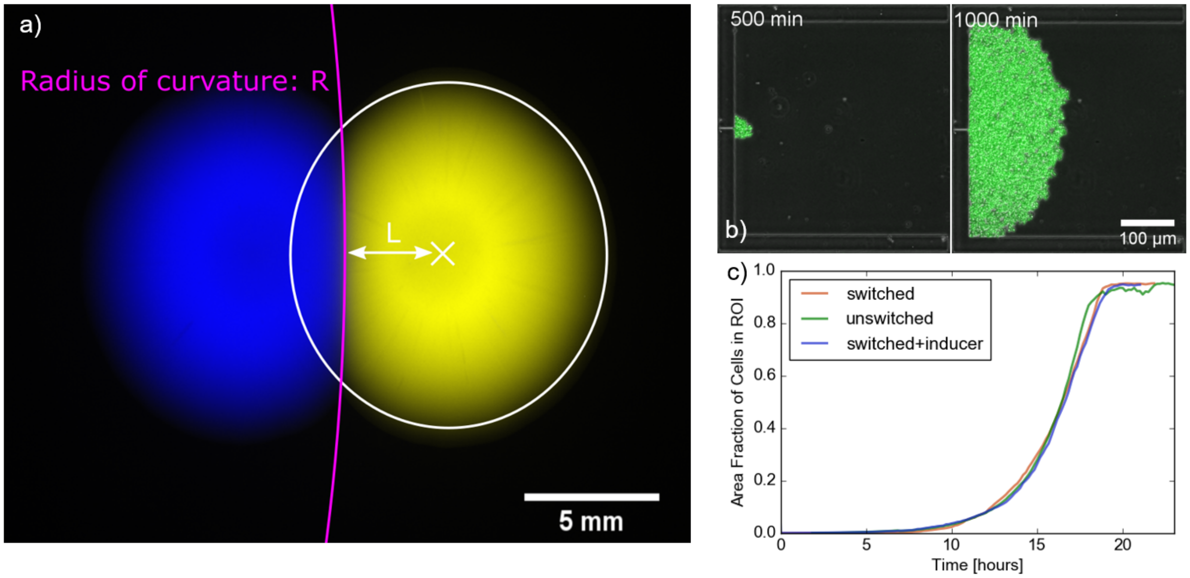


**Figure S5.** Estimation of the relative fitness between the original yJK10 strain and the color-switched yJK10 strain. (a) Colony collision experiments to estimate the fitness effect of color switching. Collisions of 12 pairs of the original yJK10 colony and the color-switched yJK10 colony were observed on YPD plates. The relative fitness between two strains was estimated at $s = 0.022 \pm 0.040$ by the formula $s = L/R$ from the equal time argument korolev2012. The lines on the figure are illustrations of the concept and not the actual fittings. (b) Population expansion experiments in microfluidics. 1-3 cells were initially trapped in the microfluidic chamber, and the growth of the population was observed for the original yJK10 strain (with YPD) and the color-switched yJK10 strain (with YPD and YPD + $\beta$-estradiol). (c) The exponential fitting of the growth curves gives us the estimation of the relative fitness of the color-switched strain to the original strain: $s = 0.019$ (YPD) and $s = -0.020$ (YPD + $\beta$-estradiol).

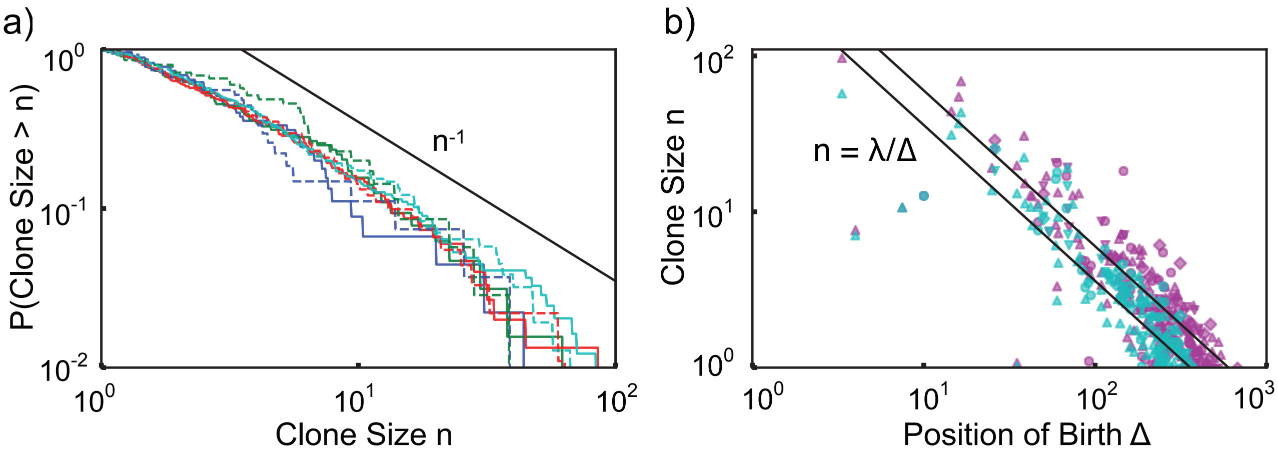


**Figure S6.** Proportion of color-switched cells whose final clone size is greater than n, where area is used as a proxy for clone size. The different lines indicate experimental replicas with respectively 45 (blue), 64 (green), 150 (red), 245 (cyan) mutant clones. The solid lines correspond to the chamber depth of $\lambda = 500$ µm as used in Fig. 2c and the dashed lines correspond to clones imaged at a distance $\lambda = 300$ µm, mimicking the clones we would expect to see in a shorter chamber. (d) Relationship between final clone size and distance from the front at which such clone arose. Purple point correspond to $\lambda = 500$ µm and cyan points correspond to $\lambda = 300$ µm. The different point types indicate experimental replicas with respectively 45 (diamonds), 64 (upside down triangles), 150 (circles), 245 (rightside up triangles) mutant clones. The black line corresponds to $\lambda/\Delta$, where $\lambda$ is the size of the chamber and $\Delta$ is the distance from the front.

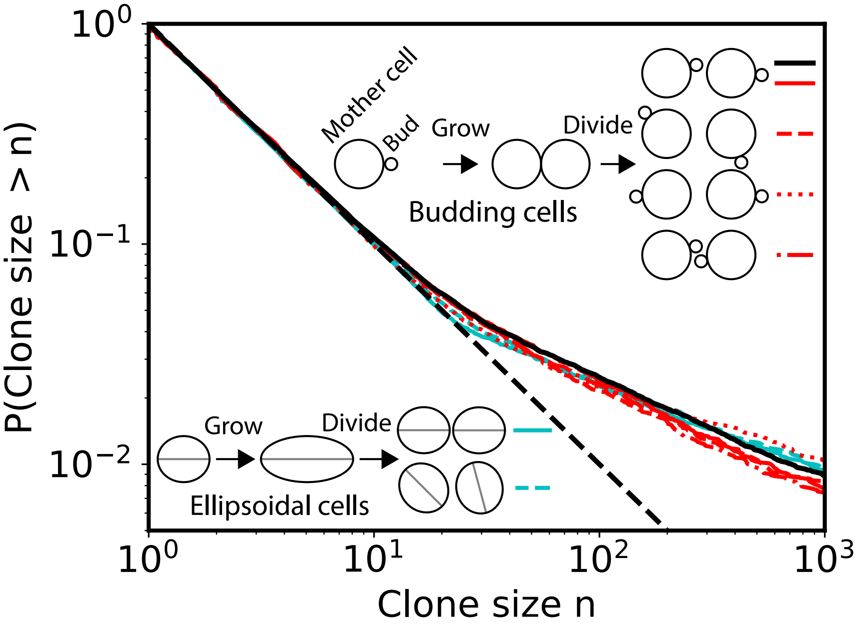


**Figure S7.** Clone size distributions for neutral mutations in mechanical simulations of ellipse-shaped and budding cells with different division rules (growth layer depth $\lambda = 14$ cell widths). The ellipse-shaped cells in these simulations have aspect ratio $= 1$ at birth and grow to aspect ratio $= 2$. These simulations use conjugate gradient energy minimization (see Gniewek2019) rather than overdamped molecular dynamics as used in the main text (Fig. 3). Ellipse data is shown in cyan, budding data is shown in red, and budding data from the main text (Fig. 3b) is shown in solid black for reference. The dashed black line shows the $1/n$ prediction. We compare four different rules for assigning the orientations after division, including the case where cells retain the orientation of their mothers (solid black/cyan/red lines), are assigned random orientations (dashed red/cyan lines), exhibit polar budding with new buds facing outward (dotted red line), and exhibit axial budding with new buds facing inward (dot-dashed red line).

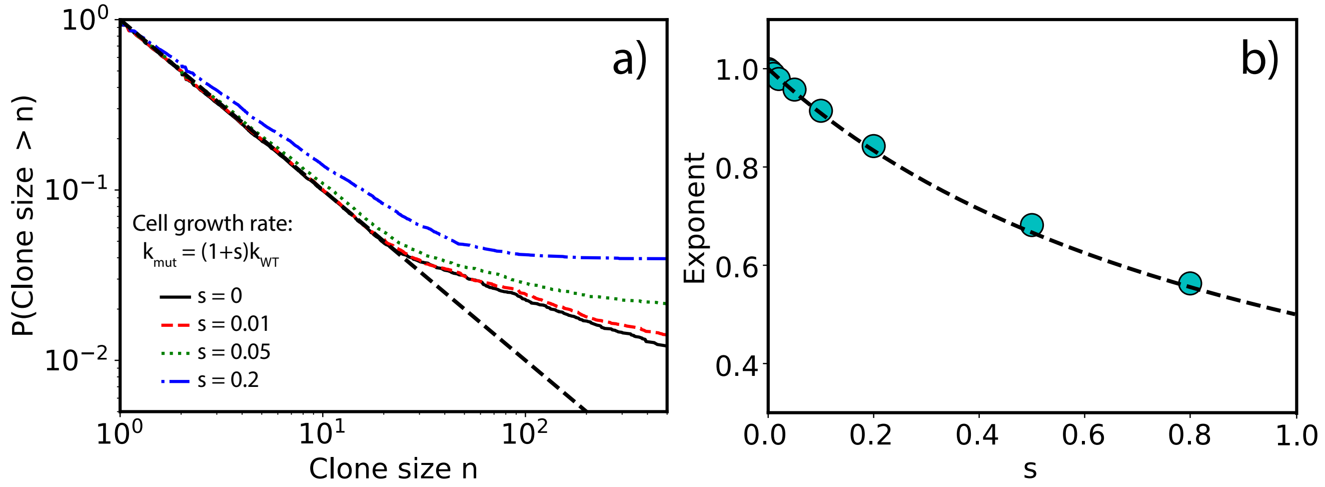


**Figure S8.** Clone size distributions for advantageous mutations in mechanical simulations with a uniform growth layer depth of $\lambda = 14$ cell widths. (a) Distributions for selective advantages ($s = k_{\mathrm{mut}}/k_{\mathrm{WT}} - 1$) of $s = 0$ (solid black), $s = 0.01$ (dashed red), $s = 0.05$ (dotted green), and $s = 0.2$ (dash-dotted blue). The dashed black line show the $1/n$ prediction. (b) The small-$n$ power-law exponent (cyan points), found in the range $n < 10$, compared to the predicted value $P(\text{Clone size} > n) \propto n^{-1/(1+s)}$ (dashed black line). For these simulations, we used ellipse-shaped cell simulations where cells have aspect ratio $= 1$ at birth and grow to aspect ratio $= 2$. These simulations use conjugate gradient energy minimization for population dynamics.

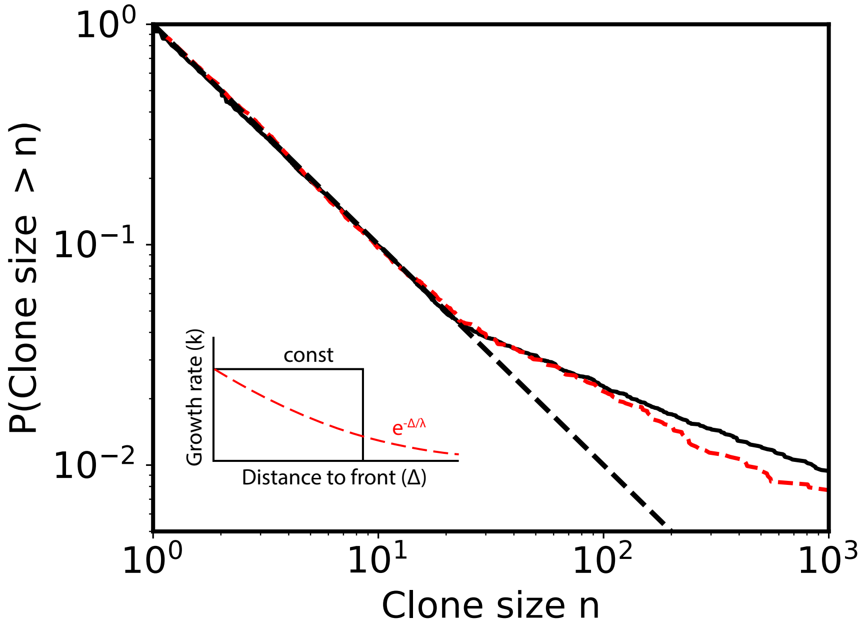


**Figure S9.** Clone size distributions for neutral mutations in mechanical simulations with a uniform growth layer (solid black line) and a growth layer profile where cellular growth rate decreases exponentially with distance to front (dashed red line). The dashed black line show the $a/n$ prediction. Both simulations have a characteristic growth layer depth of $\lambda = 14$ cell widths. For the uniform growth layer, the growth rate $k = k_0$ for $\Delta < \lambda$ and $k = 0$ for $\Delta > \lambda$, where $\Delta$ is the distance to the colony front. For the uniform growth layer, the growth rate $k = k_0 \exp -\Delta/\lambda$ for $\Delta < \lambda_{\text{cut}}$ and $k = 0$ for $\Delta > \lambda_{\text{cut}}$, where we used a cut-off distance of $\lambda_{\text{cut}} = 40$ cell widths. For these simulations, we used ellipse-shaped cell simulations where cells have aspect ratio $= 1$ at birth and grow to aspect ratio $= 2$. These simulations use conjugate gradient energy minimization for population dynamics.
